# The pneumococcal bacteriocin streptococcin B is produced as part of the early competence cascade and promotes intraspecies competition

**DOI:** 10.1101/2024.10.01.616058

**Authors:** JD Richardson, Emily Guo, Ryan M Wyllie, Paul Jensen, Suzanne Dawid

## Abstract

*Streptococcus pneumoniae* is an important human pathogen that normally resides in the human nasopharynx. Competence mediated bacteriocin expression by *S pneumoniae* plays a major role in both the establishment and persistence of colonization on this polymicrobial surface. Over 20 distinct bacteriocin loci have been identified in pneumococcal genomes, but only a small number have been characterized phenotypically. In this work, we demonstrate that 3/4th of *S. pneumoniae* strains contain a highly conserved *scb* locus that encodes an active lactococcin 972-like bacteriocin called Streptococcin B. In these backgrounds, the *scbABC* locus is part of the early competence cascade due to a ComE binding site in the promoter region. Streptococcin B producing strains target both members of the population that have failed to activate competence and the 25% of the population that carry a naturally occurring deletion of the ComE binding site and the functional bacteriocin gene. The ComR-type regulator found directly upstream of the *scb* locus in *S. pneumoniae* strains can activate *scb* expression independent of the presence of the ComE binding site but only when stimulated by a peptide that is encoded in the *scb* locus of *S. pseudopneumoniae*, a closely related bacterium which also inhabits the human nasopharynx. Given the co-regulation with competence and the phenotypic confirmation of activity, Streptococcin B represents a previously unrecognized fratricide effector that gives producing strains an additional advantage over the naturally occurring deleted strains during colonization.

**Importance:** *Streptococcus pneumoniae* is a common cause of pneumonia, meningitis, sinusitis and otitis media. In order to successfully colonize humans, a pre-requisite to the development of invasive disease, *S. pneumoniae* must compete with other bacterial inhabitants of the nasal surface for space and nutrients. Bacteriocins are small antimicrobial peptides produced by bacteria that typically target neighboring bacteria by disruption of the cell surface. *S. pnuemoniae* encodes a large number of potential bacteriocin, but, for most, their role in competitive interactions has not been defined. This work demonstrates that isolates that produce the bacteriocin Streptococcin B have an advantage over non producers. These observations contribute to our understanding of the competitive interactions that precede development of *S. pneumoniae* disease.

---

*Streptococcus pneumoniae* resides in the human nasopharynx where it must compete with other members of the nasopharyngeal community for nutrients and space. This competition is in part carried out by bacteriocins which are small antimicrobial peptides that target local, non-self, bacteria[1-3]. The pneumococcal genome contains a variety of loci encoding bacteriocins, some are found in all pneumococcal genomes while others are only found in a subset [4]. The expression of many of the pneumococcal bacteriocins is tied at the regulatory level to the competence state. For example, the universally conserved CibAB bacteriocins play a role in competence mediated fratricide by targeting members of the pneumococcal population that have failed to trigger competence and therefore are not producing the immunity protein CibC [5]. CibAB expression has been shown to play a role in protecting a resident population of *S pneumoniae* against invasion during the early stages of colonization in the mouse model [6]. The production of *blp* encoded bacteriocins is indirectly tied to competence via secretion of the *blp* inducing peptide, BlpC, by the ComAB transporter [3, 7, 8]. The *blp* locus is found in all genomes but, unlike the *cibABC* locus, the bacteriocin/immunity content is variable from strain to strain allowing for intra-species competition that is independent of the activation state of the neighboring strain. We have shown that the *blp* locus a plays a role in intraspecies competition in vivo, especially in the subset of isolates that encode a dedicated BlpAB transporter [2, 3, 7, 9, 10]. Isolates with an intact BlpAB transporter can both activate the *blp* locus independently of competence and are characterized by more permissive activation of competence due to the secretion of competence stimulated peptide (CSP) by BlpAB. The variable content and expression of bacteriocins in any individual isolate plays a significant role in dictating the outcome of any interaction with the resident community. Of the remaining twenty or more described bacteriocin loci found in pneumococcal genomes, only the *blp* bacteriocins, CibAB and the locus encoding Pneumolancidin K have phenotypic confirmation of their inhibitory activity [2, 5, 10-12].

In this work, we characterize the regulation, function and distribution of one of the five known *S pneumoniae* encoded lactococcin 972-like bacteriocins called Streptococcin B. We show that a version of the *scb* locus encoding Streptococcin B is found in nearly all pneumococcal genomes and, in a majority of isolates, is a previously unrecognized component of the early competence regulon. A subset of strains including the common laboratory strains Tigr4, D39 and R6 lack the competence regulatory region as well as the functional bacteriocin peptide. These strains do not appear to endogenously activate the locus including the dedicated immunity proteins under any tested growth conditions. However, *scb* expression does occur in these strains via the locus-associated ComR homolog, but only when stimulated by a XIP encoded within the homologous *scb* locus of *S. pseudopneumoniae*. D39-like strains are inhibited by isolates expressing the fully functional locus in plate assays and during biofilm growth under competence permissive conditions. Streptococcin B is an additional member of the previously characterized fratricide effectors, allowing for inhibition of both non-competent and Streptococcin B negative neighbors.

## Results

### Characterization of the variability of the *scb* locus in pneumococcal genomes

In the context of understanding the contribution of bacteriocin loci controlled by peptide pheromone-based quorum sensing systems, we focused on the *scb* locus previously described by Rezaei Javan et al [4]. The *scb* locus was predicted to encode a lactococcin 972-like bacteriocin called Streptococcin B in addition to ScbB and ScbC which are predicted immunity proteins. The locus was found directly downstream of a previously uncharacterized ComR-type transcriptional regulator suggesting a role in regulation of the locus. Rezaei Javan et al found the *scb* locus in all 571 genomes queried but with variable completeness. Of the 571 genomes, 414 (72.5%) were found to have a complete *scbABC* operon while 157 (27.5%) only encoded the immunity proteins ScbBC (Fig 1A). To better understand the contribution of this locus to pneumococcal competition, we created a series of reporter strains where the *scbBC* promoter region from D39 or the *scbABC* promoter region from the SP9BS68 strain were joined to a luciferase gene and cloned this into the transcriptionally silent CEP locus in a D39 background created either with or without an intact *blpA* gene. We additionally created a reporter that incorporated the HiBiT tag into the C terminal end of the Streptococcin B peptide with the luciferase gene placed in operon structure after the scbA_HiBiT_ gene to assess secretion of Streptococcin B. These reporter strains were grown in chemically defined media at pH 6.8 and 7.4. The D39 derived *scbBC* promoter did not appear to activate under any of these conditions, but the SP9BS68 *scbABC* promoter activated at pH 7.4 in a *blpA* deficient background but at both pH 6.8 and 7.4 in the background of a *blpA* sufficient strain. (Fig 1B).

**Figure 1.**
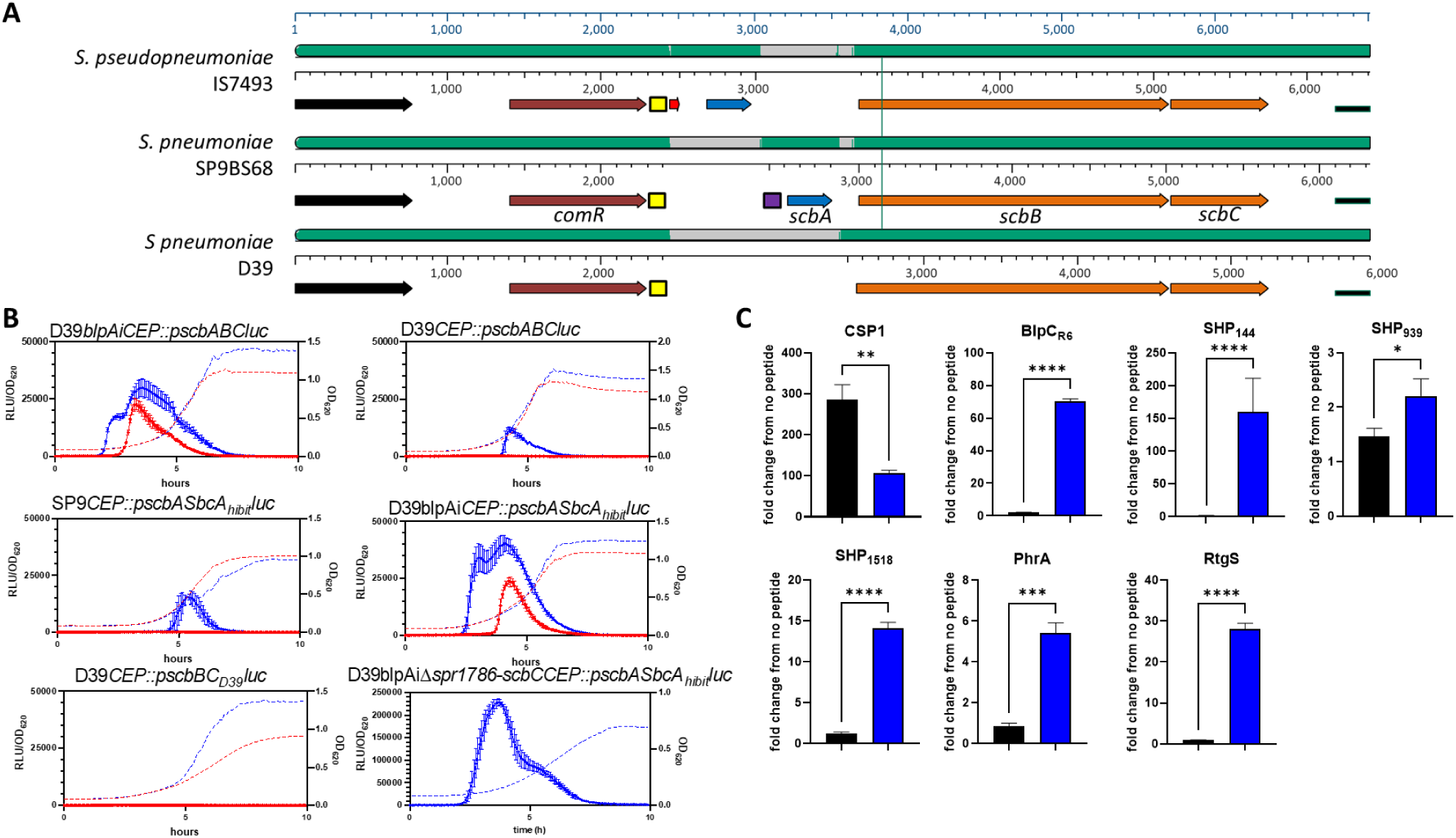
Diagrammatic representation of the *scb* loci and activation characteristics. A. Mauve alignment of the D39, SP9BS68 and *Streptococcus pseudopneumoniae scb* loci. ORFs are depicted as arrows, binding sites as boxes. The ComR encoding gene referred to as SPD_1876 in D39 is shown in brown, the *scbA* gene encoding the functional bacteriocin is shown in blue and the genes encoding the ScbB and C immunity proteins are shown in orange. The putative SHP ORF identified in *S. pseudopneumoniae* is shown as a red arrow. The ComE binding site and ComR Binding sites are shown as purple or yellow boxes, respectively. Areas of homology to the *S pseudopneumoniae* sequence are shown as green shading, areas lacking homology are grey. B. Promoter activity of the various *sbc* promoter-luciferase fusions during growth in CDM at pH 7.4 (blue) and pH 6.8 (red). RLU/OD_620_ is shown on the left Y axis and represented by solid lines, while OD_620_ alone is shown on the right Y axis and represented by dotted lines. C. Promoter activation of either the D39*CEP::pscbSbcAHiBiTluc* strain (black bars) or the peptide specific reporter strain (blue bars) 45 minutes following addition of 250 ng/mL of peptide. Activation is shown as fold change from the same reporter with vehicle only addition. All reporters were created in the D39 background (*blpA* deficient).

### Activation of the *scbABC* locus by CSP and *S. pseudopneumoniae* XIP

We initially attempted to activate the *scbABC* locus using 250 ng/mL of a 10 AA version of a hypothesized XIP encoded in both the *S. pneumoniae scbBC* and *scbABC* loci (SYQNSSKVRL). However, addition of this peptide did not activate either reporter construct (not shown). To determine if other, previously described, peptide signals were involved in *scb* control, 250ng/mL of the peptides SHP143, SHP939, SHP1518, PhrA, RtgS, BlpC2 and CSP1 were used to induce expression of the *scbABC* reporter strain. Control reporters were used for each peptide in the D39 background to verify induction of peptide responsive promoters (Fig 1C). In each case, peptides induced their respective control reporters but only CSP1 induced the expression of both the control reporter and the SP9BS68 derived *scbABC* promoter. Addition of CSP1 did not induce expression of the D39 derived *scbBC* promoter (not shown).

In surveying other streptococcal species for the presence of *scbA* homologues, we identified a *Streptococcus pseudopneumoniae* strain (SPPN IS7493) with a highly homologous *scbABC* locus. The *S. pseudopneumoniae scbABC* locus also contained an upstream ComR homolog. By comparing the nucleotide sequence downstream of the ComR homologs and upstream of the *scbBC* or *scbABC* locus, we identified a canonical ComRS promoter motif– taag**gacat**tg(**a/g)tgtc**cttg-n_20_-**tataat** – in *S. pseudopneumoniae* as well as in SP9-type and D39-type strains of *S. pneumoniae*. Interestingly, only in *S. pseudopneumoniae* were we able to identify the expected cognate *comS* gene downstream of the promoter motif. The *S. pseudopneumoniae comS* gene encodes a short 19-AA peptide with the canonical Type II XIP double tryptophan motif: MFGFIMFLTYFSF**<u>GDWWHG</u>**. Based on the well characterized ComRS systems in *Streptococcus mutans*, we predicted that the mature XIP sequence would be the final six C-terminal amino acids. This putative XIP was synthesized and tested against the reporter strains. The six AA XIP peptide successfully activated both the D39 *scbBC* and SP9 *scbABC* promoters. Consistent with previously characterized ComRS systems, this stimulation is dependent on an intact AmiCDE importer. No stimulation was noted in a ComR deletion background (Fig 2), suggesting that this regulator interacts with the peptide. A survey of the pneumococcal SP9 and D39 genomes does not reveal any region that would be predicted to encode this peptide. Alignment of the *comR*-*scbABC* intergenic region from the *S. pseudopneumoniae* strain (Figure 1A) demonstrates that the region encoding the XIP peptide was likely exchanged for the ComE binding site at some point early in pneumococcal evolution.

**Figure 2.**
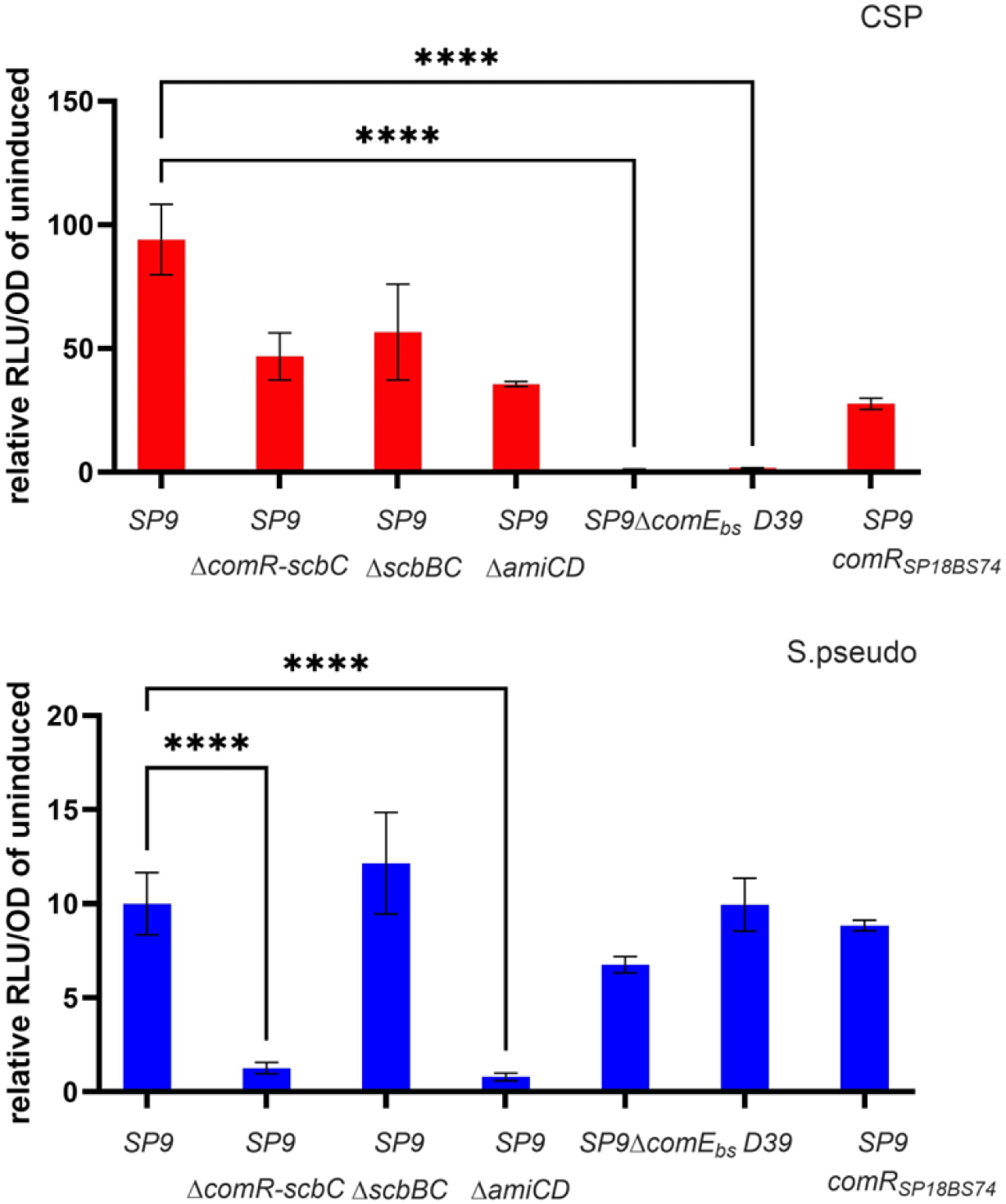
Role of the *scb* locus in regulation of Streptococcin B production. D39 strains with the marked p*scb* reporters were grown in CDM 6.8 and induced with 250ng/mL of CSP1 (red), S pseudopneumoniae XIP peptide (blue) or vehicle control at 37°C. Luminescence/OD values were determined after 30 minutes of incubation and compared with the no peptide control for each reporter. Each reporter strain is designated on the graph by the origin of the promoter content on top (SP9, SP9 with ComE binding site deletion or D39) and any additional chromosomal mutation below. Reporter strains were tested in the D39 *blpA* intact background and compared by one way ANOVA for significance.

### Verification of a ComE binding site in the *scbABC* promoter

Given the early, robust stimulation of the *scbABC* locus by CSP1, we examined the promoter for a ComE or ComX binding site. A classic ComE binding site (catttcagggtgctattttgacaatttag) was identified in the *scbABC* promoter 115bp upstream of the start codon of *scbA*. This sequence was not found in the D39 derived *scbBC* promoter region (Fig 1A). Alignment of these two *scb* loci demonstrated that the deletion that resulted in loss of the *scbA* gene in D39 also included the region of the promoter that contains the ComE binding site (Fig 1A). To demonstrate the role of the ComE binding site in CSP-mediated regulation of the *scbABC* locus, we created a reporter strain where just the ComE binding site was deleted. As expected, this strain no longer activated luciferase expression in response to addition to CSP although activation with the *S. pseudopneumoniae* XIP was retained (Fig 2). To determine if the upstream ComR homolog played a role in competence-mediated control of the *scb* locus, we created a deletion of the entire *scb* region including the ComR regulator gene in our existing reporter backgrounds. These reporters had identical *scbABC* regulation compared with intact backgrounds when allowed to induce naturally (fig 1B) or when stimulated with CSP (Fig 2), indicating that CSP-induced activation of *scb* is independent of ComR. Retained natural activation of *scbABC* the *comR* deletion strains additionally suggests that, unlike what is found in other Strep species, *comR* is not required for activation of competence.

The D39 *scbBC* region was included in the transcriptome analysis done by Aprianto et al [13]. In their accompanying searchable database, Pneumoexpress (https://veeninglab.com/pneumoexpress-app/), both *scbB* (SPV_1785) and *scbC* (SPV_1784) show 2-3 fold increases in average transcript following either 30 or 60 minutes of exposure to live A549 cells. In an attempt to determine if the D39 locus might induce in response to factors secreted by epithelial cells, we reproduced the cell exposure as described by Aprianto using the D39 strain with the p*scbBC* reporter and compared activation of the reporter with cells in media alone. This strain did not show any evidence of activation following exposure to epithelial cells (Fig S1)

### Contribution of *blpA* status to the activation of the *scbABC* promoter

The status of the bacteriocin encoding *blp* locus can have a substantial impact on competence development. Strains that encode an intact BlpAB transporter induce competence earlier in the growth phase and under typically non permissive conditions. This is due to secretion of CSP through the BlpAB transporter, augmenting its accumulation [3, 14]. To determine how BlpA intact strains induce the intact *scbABC* locus differently than those that lack a functional BlpAB, we allowed identical *scb* reporter strains created in otherwise isogenic *blpAB* intact and Δ*blpA* backgrounds to grow in CDM at pH 6.8 and 7.4. Under these conditions, only *blpAB* intact strains activated uniformly at pH 6.8 and 7.4 (Fig 3). The Δ*blpA* strain activated only ∼60% of wells at pH 7.4 and at higher ODs compared with *blpAB* intact strain at the same pH.

**Figure 3.**
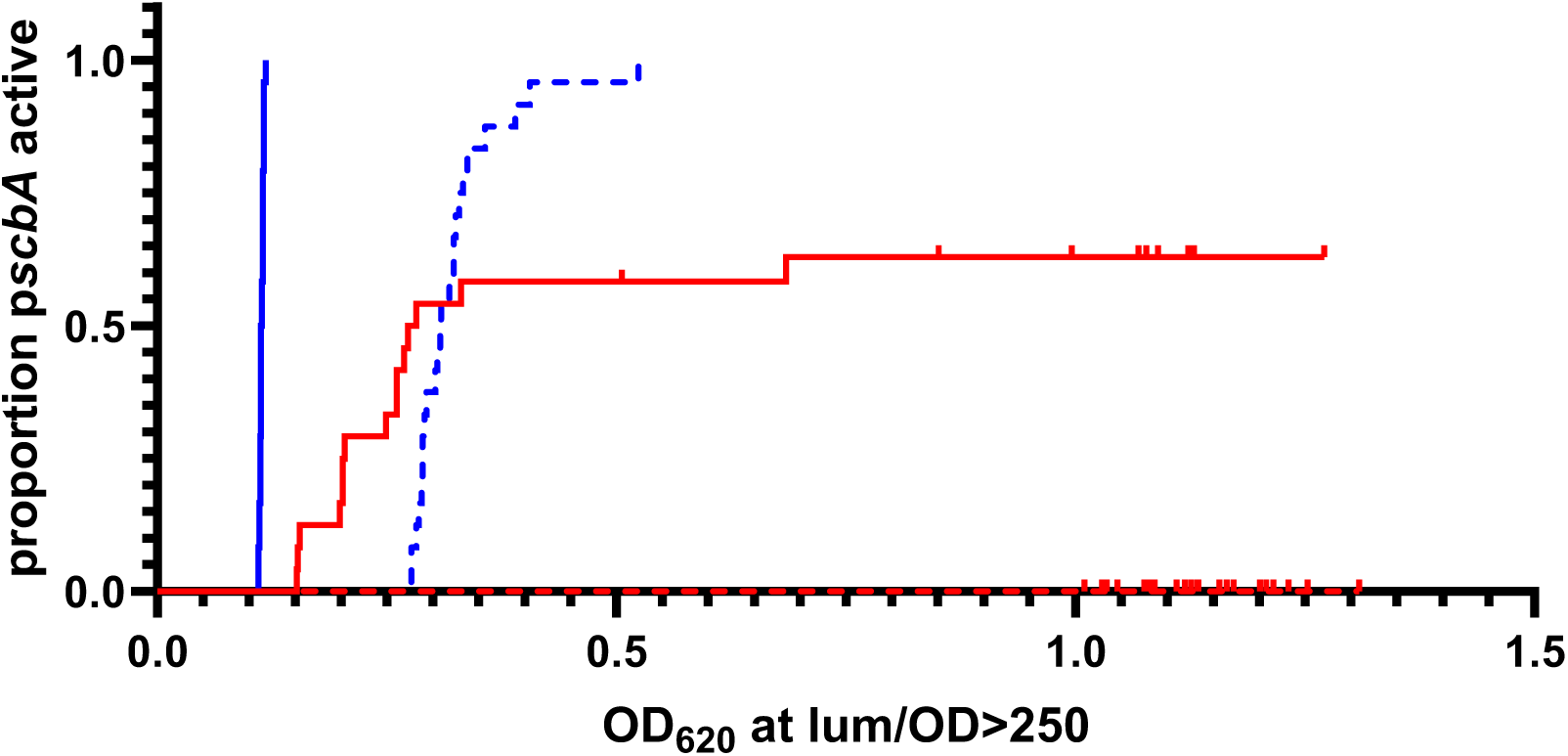
BlpA intact strains activate *scbABC* promoter at low and high pH. The *scbABC* reporter was grown in CDM at pH 6.8 (red) and 7.4 (blue) in the *blpA* intact (solid) and frame shifted (dashed) backgrounds. 24 wells were used to assess each strain/condition in a 96 well plate. The OD was noted when an individual strain exceeded the threshold of 250 RLU/OD_620_. Any wells that did not reach this threshold were right corrected.

### Distribution of the *com* activated *scbABC* locus in a defined collection of pneumococcal genomes

We surveyed the 617 genomes analyzed by Mitchell et al consisting of 15 different Strain Clusters (SC) to determine the distribution of competence regulated *scb* loci (Fig 4). Consistent with the findings of Rezaei Javan et al, we found that 150 genomes (24%) had the D39-like *scbBC* locus and 466 (76%) had the SP9-like *scbABC* locus. All genomes were predicted to encode a ComR regulator upstream of *scb*. Eighty-four of 617 genomes encode an allelic variant of the more common SPR_1786 *comR*. This variant has 73% similarity and 57% identity to SPR_1786 with the majority of the identity in the N terminal 103 AA (101/103 identity). This variant was moved into the D39 reporter background replacing the native comR gene. This reporter was shown to also respond to *S pseudopneumoniae* peptide (Fig 2). All 466 SP9-like loci had the identified ComE binding site. The full length *scbABC* locus was found in 11 of 15 SC while the *scbBC* locus was found in 7 of 15 SC. There was no absolute correlation between CSP type and the competence regulated version of the locus; CSP1 encoding strains made up 358/466 (77%) of the *sbcABC* strains compared with 396/617 (64%) total strains. Considering the difference in activation of the *sbcABC* locus noted in *blpA* intact vs disrupted backgrounds, the distribution of BlpA expressing strains was examined. Of the 179 genomes with a predicted intact *blpA* gene, 139 (77%) are predicted to encode *scbA*, similar to the distribution in the whole population.

**Figure 4.**
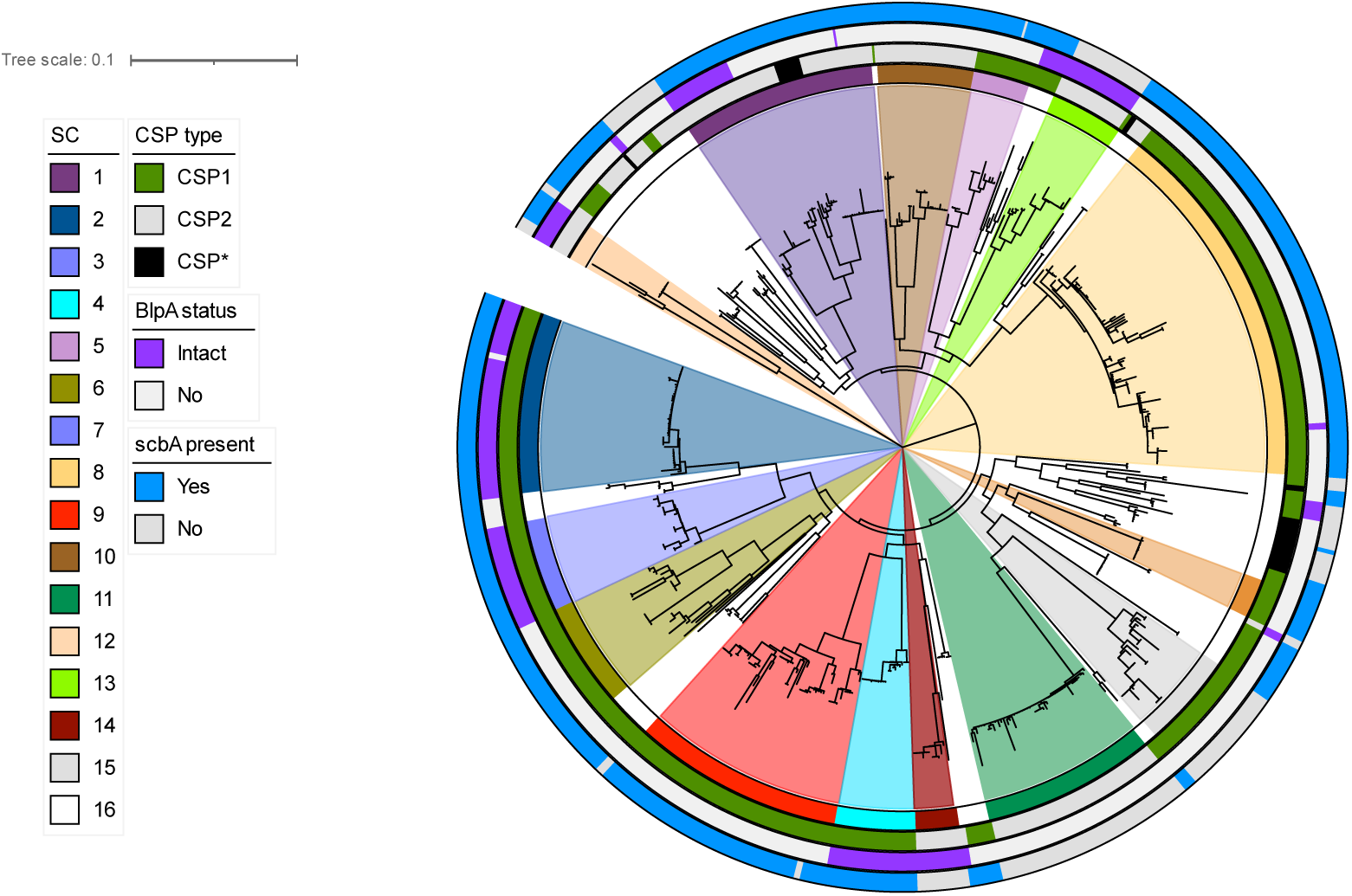
Distribution of the SP9 and D39 like version of the *scbA* locus in the Boston Isolate collection. Genomes were searched for SP9 *scbA* sequences in addition to *blpA* and *comC* sequences using the PubMLST website. SC designations assigned by Mitchell et al [14] are shown in colored wedges, *blpA*, *scb* and CSP content is noted in outside rings. The tree was recreated using the newick file provided in Mitchell et al. using iTOL.

### Streptococcin B is not secreted by BlpAB or ComAB

Lactococcin 972-like bacteriocins are thought to be secreted by the Sec system and do not require a specialized processing and secretion system like the Cib and Blp bacteriocins. Given the regulatory connection to competence stimulation, we investigated whether the ComAB transporter was required for Streptococcin B secretion. Using the reporter expressing a HiBiT tagged version of Streptococcin B, we assayed culture media for HiBiT tag secretion following CSP secretion in strains expressing or lacking the ComAB transporter. Of note, the D39 background naturally lacks other potential bacteriocin transporters including RtgAB and BlpAB. HiBiT tagged Streptococcin B was secreted into the media following CSP stimulation in both ComAB containing and lacking backgrounds in support of secretion via the Sec system (Fig 5A).

**Figure 5.**
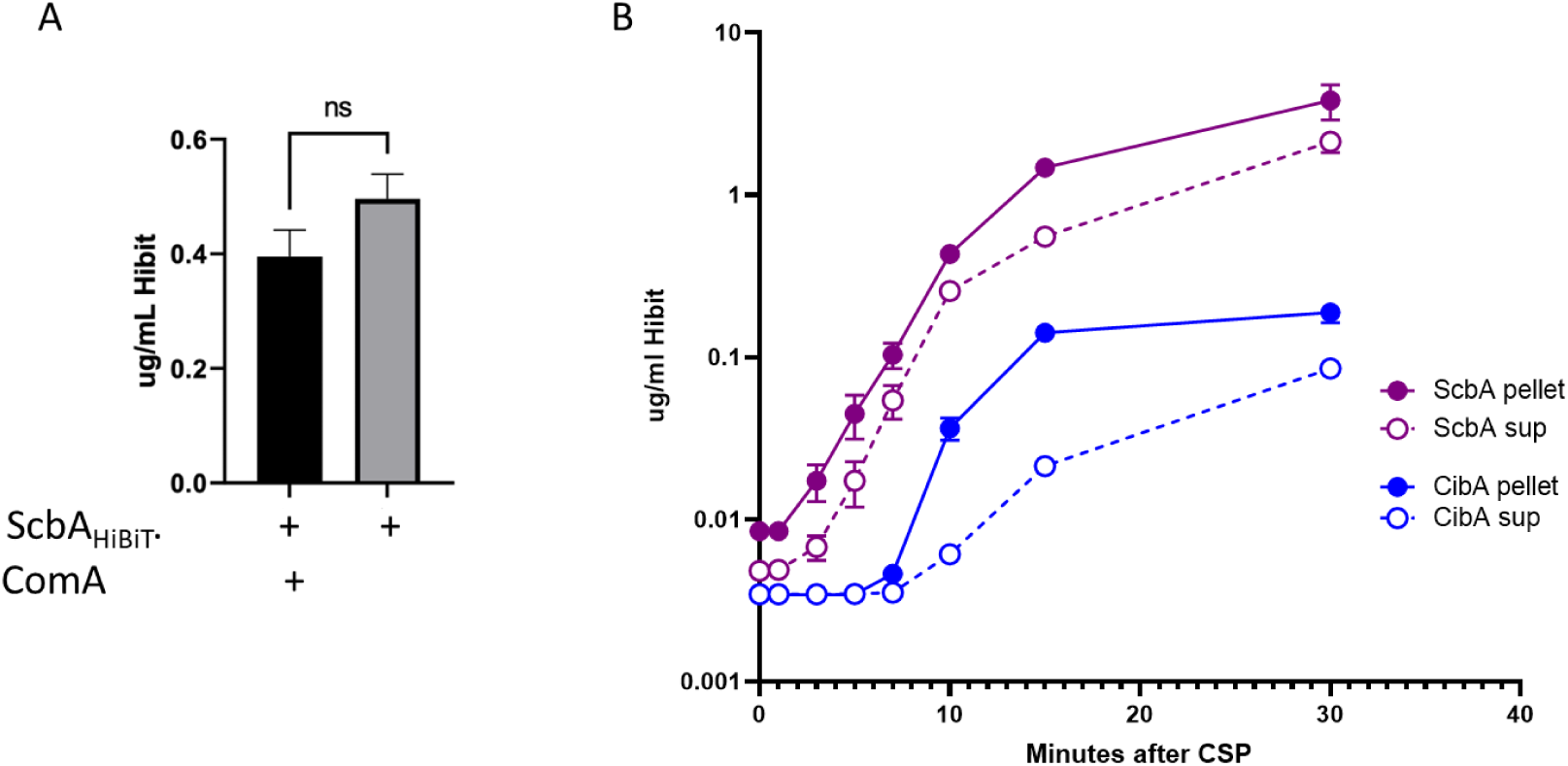
Secretion of ScbAHiBiT is independent of BlpAB and ComAB and occurs early in competence. A. D39 reporters with ScbAHibiT fusions were grown in CDM and induced with CSP in a *comA* sufficient and deleted background. Following 1 hour of stimulation, supernatants were assessed for the concentration of the HiBiT tag. Quantification was determined using dilutions of the idealized HiBiT peptide. B. ScbHiBiT or CibAHiBiT reporter strains were stimulated with CSP at time = 0. Samples were removed at 0,2,5,7, 10, 15 and 30 minutes, centrifuged and supernatants and pellets were placed on ice. Resultant samples were assayed for HiBiT concentration using a standard curve generated with the HiBiT standard peptide.

### Streptococcin B is produced earlier and is highly secreted compared with CibA

To compare the secretion kinetics of Streptococcin B with that of the fratricide effector bacteriocin, CibA, which is encoded by a late competence gene and dependent on ComAB secretion, we performed a time course experiment using otherwise isogenic strains expressing either CibAHiBiT or StreptococcinBHiBiT. StreptococcinB HiBiT was detectible in the media in as little as 3 minutes post stimulation while CibA secretion was delayed in comparison, detected at between 7 and 10 minutes following CSP stimulation (Figure 5B). A larger percentage of HiBiT tagged Cib remained cell associated compared with Streptococcin B. Although we cannot exclude differential effects on secretion brought about by the addition of the HiBiT tag, these findings are consistent with Streptococcin B being more efficiently secreted from the cell. These findings suggest that StreptococcinB plays a significant and early role in competence-mediated fratricide.

### Streptococcin B inhibits *scbBC* expressing strains in plates and biofilms

To determine if Streptococcin B production produces detectible inhibition in vitro, we performed overlay assays using the SP9BS68 wildtype strain, D39, and D39 with the *sbcBC* locus replaced with the SP9 *scbABC* locus. No inhibition was noted in the absence of CSP addition, however, when CSP was added to the stabbed strain, SP9 and D39 expressing the SP9 version of *scbABC* both showed zones of inhibition when tested against D39 or D39*ΔscbABC* (Fig 6A). Zones of inhibition could still be appreciated in a Δ*lytA*, Δ*cibAB* and Δ*cbpD* background demonstrating that the inhibition noted was not the combined result of these fratricide effectors and rather related to expression of the *scbABC* locus alone.

**Figure 6.**
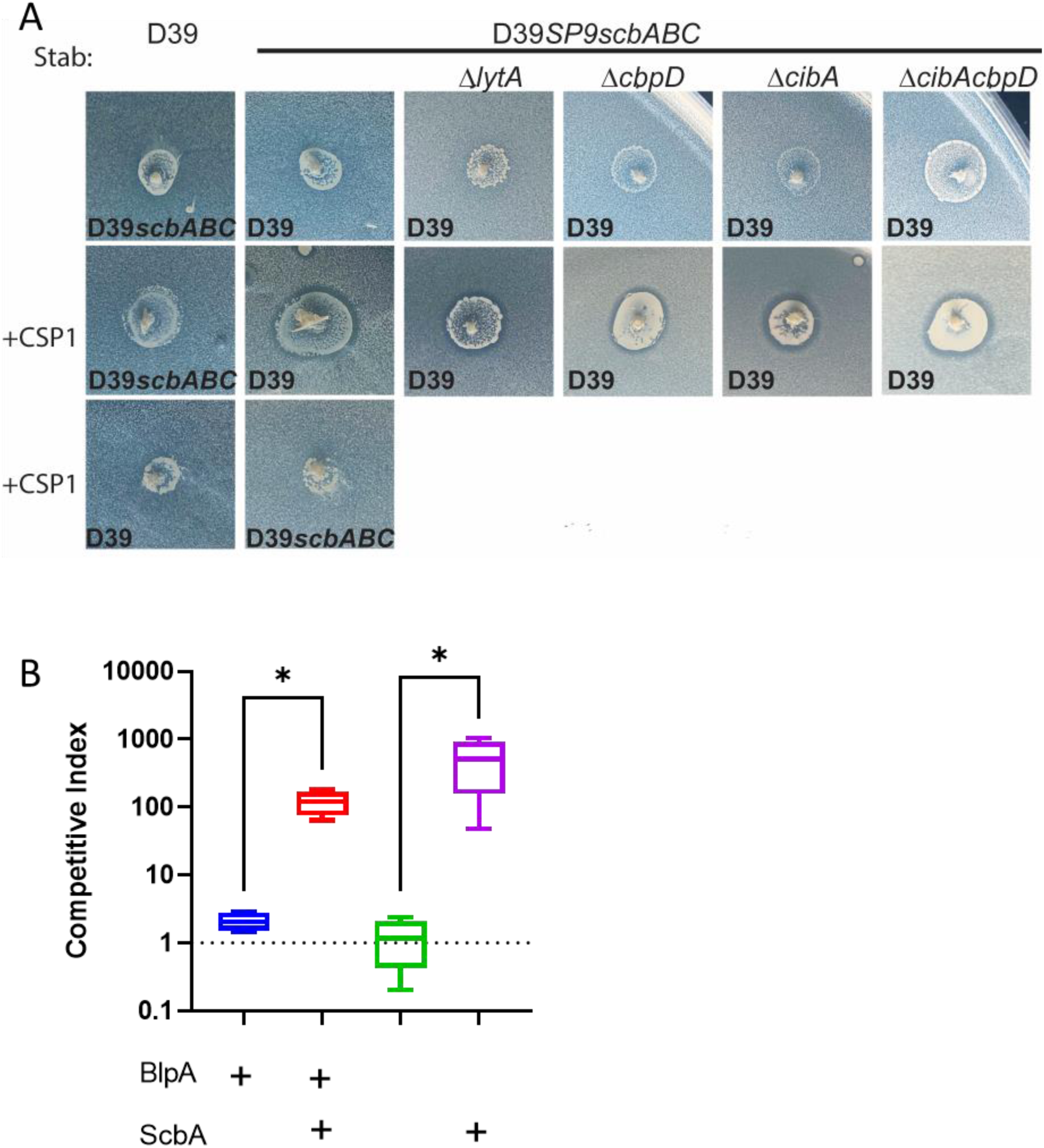
SP9 like *scbABC* expression results in phenotypic inhibition of D39 *scbBC* expressing strains in plate and biofilm assays. A. Overlay assays demonstrating the ring of inhibition noted in *scbABC* containing strains following the addition of CSP. The stabbed strain is shown above the photos and the overlay lawn strain is marked on each individual photo. B. Competitive index comparing the spectinomycin resistant strain with either SP9 like *scbABC* or D39 like *scbBC* with a kanamycin/streptomycin otherwise matched competitor strain following overnight incubation in CDM 7.4 over a fixed epithelial cell monolayer. * P<0.05 by Mann Whitney.

To determine if inhibition of *scbBC* expressing strains could be noted during growth in a biofilm, we created the *scb* variants in previously described D39 derivatives in which the inactive *blp* locus was replaced with an active *blp* locus either with or without a frame shift mutation in *blpA*. We have previously described the behavior of these strains in biofilm competition [7]. The competitor strains carry a spectinomycin cassette directly upstream of the *blp* locus and the sensitive D39 strain carries streptomycin and kanamycin resistance markers. Because all backgrounds are matched at the *blp* locus, no *blp* mediated competition is appreciated in these pairings. Equal inocula of the competitor and dual resistant strains were added to wells with a fixed epithelial substratum and allowed to establish overnight before plating. In static biofilm conditions, both strains carrying the SP9-like *sbcABC* had a competitive advantage compared with the D39-like *scbBC* in both the blpA sufficient and deficient backgrounds (Fig 6B).

## Discussion

The pneumococcal population is characterized by significant diversity in genome content. There are at least 23 distinct bacteriocin or bacteriocin-like loci that are encoded in the pneumococcal chromosome but only a handful of these have been characterized at the phenotypic level [4]. Streptococcin B is in the family of lactococcin 972-like bacteriocins. Unlike traditional alpha helical bacteriocins, this family forms a beta sandwich that is not predicted to participate in pore formation [15]. These bacteriocins have been shown to function by interacting with lipid II and inhibiting septum formation [16-18]. The lipid II binding site appears to be separate from that for nisin. These bacteriocins are classically encoded in a three gene operon that consists of a gene encoding the structural bacteriocin followed by two genes encoding immunity proteins.

Bacteriocins are often controlled by a quorum sensing system in the immediate genomic vicinity that restricts production to situations where the bacteriocin is likely to accumulate. Many classes of peptide-based quorum sensing systems have been described in *S. pneumoniae*, including the Tpr-Phr system and multiple Rgg-Shp class systems [4, 19]. Here, we report for the first time the presence of a ComRS-type system in *S. pneumoniae*. ComRS systems were first described in *S. mutans* and *S. thermophilus* but have since been shown to be widespread throughout the streptococci. These systems have been demonstrated to regulate various aspects of bacterial physiology including the development of genetic competence and the production of bacteriocins [20-23]. The ComRS system identified here in SP9 and D39-type strains of *S. pneumoniae* does not appear to be required for competence but does have the capability to regulate expression of the *scb* locus. While the pneumococcal *scb* loci do not encode the *comS* gene, we show that the downstream ComR-XIP binding sites are intact, and that ComR-dependent activation of the loci does occur upon addition of exogenous *S. pseudopneumoniae* XIP. While this could be a vestigial phenotype, the widespread conservation of the *scb*-associated ComR regulator gene, its downstream ComR-XIP box containing promoter, and the *scbBC* immunity genes could indicate the system has retained evolutionary utility independent of its link to competence. It is possible that the ability to express the *scbBC* immunity genes in response to environmental XIP signaling from co-colonizing *S. pseudopneumoniae* strains provides both SP9 and D39-type *S. pneumoniae* strains with protection from resulting Streptococcin B production. Should this be the case, it would represent an interesting interspecies interaction between *S. pneumoniae* and its poorly understood ecological competitor, *S. pseudopneumoniae*.

However, in most pneumococcal strains, the *scb* promoter region has gained a ComE biding site, allowing for coregulation and linking Streptococcin B production to competence. In strains without the ComE binding site endogenous activation of the locus presumably does not occur. Despite encoding intact immunity proteins, the lack of activation results in these strains being killed by Streptococcin B encoding strains under conditions of competence induction. Given the prevalence of the *scbABC* locus in the pneumococcal population in over ¾ of the genomes surveyed, this represents a previously unappreciated fratricide effector. The role of the *scbABC* locus as an early competence gene was not appreciated in early studies because the laboratory strains D39 and TIGR4, in which studies were performed, are both missing the ComE binding site. *scbABC* is produced at high amounts immediately following competence induction when compared with the kinetics of the *cibABC* operon. Additionally, because it is induced early and does not require a dedicated transporter, a greater proportion of peptide is secreted early during competence. Based on these observations, Streptococcin B production is likely to play and equivalent or greater role in fratricide compared with known fratricide effectors, in both members of the population that have not induced competence or neighboring strains that lack the ComE binding site.

The impact of an intact BlpAB transporter on non-*blp* bacteriocin mediated competition via enhanced competence activation is clearly demonstrated by this locus. BlpAB intact strains activate competence under a wider array of conditions [3, 9], thereby enhancing the expression of competence regulated competitive factors including Streptococcin B. Despite this, the combination of an SP9-like *scbABC* and and intact *blpAB* is not greater than expected in the population based on the Boston genome collection. This suggests that selective pressure is not promoting this combination despite the competitive advantage noted in vitro possibly due to offsetting energetic cost. Recent publications have tied *blp* locus activation to a founder effect during colonization, however, this phenomenon was only described in a BlpAB deficient background [1]. Because BlpAB deficient strains can only activate the *blp* locus via competence activation, these findings suggest that other competence regulated factors are also likely to play a role in early colonization fitness. Similarly, CibAB was shown to be required for blocking the new acquisition of pneumococcal isolates during early colonization using the infant mouse model [6]. This work did not address the contribution of the streptococcinB locus to blocking new acquisition but based on in vitro activity it is likely that an advantage would be noted when competing a D39-like with an SP9-like locus.

The pneumococcal genome can encode up to five distinct lactococcin 972-like loci. Streptococcin A, B and C are found in nearly all strains while Streptococcin E and D are less common [4]. The regulatory control of A,C and D are unclear but the promoters do not contain ComE binding sites. Streptococcin E, when found in a genome, replaces a different bacteriocin locus called Pneumolancidin G and based on its location would be predicted to be controlled by the TprA regulator and PhrA peptide pheromone. No function has been attributed to any of the non-B streptococcins. Based on the studies with streptococcin B, we expect that the other four loci provide a competitive advantage to expressing strains but potentially under different regulatory conditions. Some lactococcin 972-like homologues have been implicated in altering the immune response to infection. The *S. iniae* streptococcinB homologue has been shown to interfere with the respiratory burst in monocytes and preadministration of recombinant bacteriocin resulted in disseminated infection [24]. The role of Streptococcin B on the host immune response to infection has not been investigated.

## Methods

### Growth conditions and strains used in this study

All strains were derived from either D39 or SP9BS68 as described. Strains are listed in Table S1, primers are listed in Table S2, peptides are listed in Table S3. *Streptococcus pneumoniae* was grown in THY (Todd Hewitt media supplemented with 5% yeast extract) at 37°C at pH 6.8 unless otherwise stated. For transformation, cells were grown in C+Y pH 6.8 and stimulated for transformation in C + Y pH 8.0 using 500 ng CSP1/mL. Following transformation, cells were selected on tryptic soy agar supplemented with 5 μg/ml catalase and antibiotics as follows: streptomycin 100 μg/ml, kanamycin 500 μg /ml, spectinomycin 200 μg /ml or media containing 10% sucrose for counter selection of the Janus2 cassette. For natural induction assays, cells were grown in CDM+ at the indicated pH and 330 μM firefly luciferin (88294; Thermo Fisher Scientific). The media contents for CDM+ are described in [9]. All synthetic peptides were ordered from Genescript at >95% purity. All assays were performed at least in triplicate on three separate occasions, representative assays are shown.

### Strain construction

All reporter strains were constructed by PCR using a Pfusion (Termo^TM^), a high-fidelity polymerase and HiFi Cloning (NEB^TM^) and confirmed by PCR and/or sequencing. Full details of strain construction can be found in the supplemental document.

### Peptide stimulation studies

Reporter strains were grown in CDM pH 6.8 to OD620= 0.2 in the presence of 330 μM luciferin and catalase. 250 ng/mL of each peptide was added to the reporter strains at time 0. The plate was incubated in a Synergy HTX plate reader set to read absorbance at 620 nm and luminescence every 5 min at 37°C. Luminescence at t=1 hour was compared with the same strain without peptide stimulation.

For kinetic assays, strains were inoculated in CDM at the indicated pH supplemented with catalase and luciferin using thawed starter cultures grown in THY at pH 6.8 to inhibit competence development. 100 μL of starter culture was used per 5 mL of CDM. Both OD_620_ and luminescence were measure every 5 minutes during growth at 37°C and compared with blank lanes. To compare *scbABC* stimulation profiles at different pHs in the *blpA* intact and non-intact backgrounds, the point at which the luminescence/OD value exceeded 250 was identified and the OD_620_ at this point recorded and an event as described in Wang et al [9]. 16 wells per strain were assayed with each experiment. Survival curves were created using these values to mark instances of locus stimulation. Wells that did not reach this threshold were right censored.

### Peptide secretion assays

Samples for extracellular peptide quantification were read with a Synergy HTX plate reader (Biotek) in a white 96-well plate (Costar, 3917) following addition of HiBiT Extracellular Detection Reagent (Promega, N2421) at a 1:1 ratio. For endpoint assays, each sample was quantified with three technical replicates and read 5 min following reagent addition. For time-course assays, each sample was quantified with three technical replicate and read 1 min following reagent addition. Samples for intracellular peptide quantification were pelleted at 6000×g, 5 min, 4°C and resuspended in proteinase K buffer [20 mM MES pH 6.5, 20 mM MgCl2, 0.5 M sucrose, 100 µg/mL proteinase K (Fisher Scientific, BP1700)]. A 15-min incubation at 37°C removed residual extracellular peptide by proteinase K digestion. Afterwards, proteinase K was inactivated by addition of 1 mM phenylmethanesulfonyl fluoride (Calbiochem, 7110). Cells were lysed by addition of 1% Triton X-100 followed by incubation at room temperature for 15 min. The lysed samples were then mixed with HiBiT Extracellular Detection Reagent at a 1:1 ratio in a white 96-well plate and read with a Synergy HTX plate reader. Each sample was quantified with three technical replicates using dilutions of the HiBiT tag peptide alone to create a standard curve.

### Plate overlay assays

TSA plates supplemented with catalase were stabbed with a pipet tip dipped in a starter culture of the indicated strain grown in THY to OD_620_= 0.4 and frozen in 15% glycerol. The stabs were allowed to grow for 6 hours and 500ng of CSP1 was added to each stab in 5 μl. This was allowed to dry and plates were incubated for an additional hour at 37C. A soft agar overlay of the overlay strain was applied by adding 100 μl of a thawed starter culture and catalase to 5 mL of TS plus 2mL of 55°C TSA and carefully applying this over the top of the plate. Plates were read for inHiBiTion the following day.

### Biofilm assays

H292 cells were grown to confluence on 4 chamber slides, fixed in formalin and then rinsed with 4 changes of PBS. Fixed cells were inoculated with plate grown strains resuspended to an OD_620_ of 0.050 in CDM 7.4. Strains were inoculated with or without competition into individual wells and incubated at 34°C in 5% CO_2_. Half of the media volume was removed after 6 hours of incubation and replaced with fresh media. Media was removed from biofilms after overnight incubation and the bacteria were harvested in 200 μl PBS using a small cell scraper. Dilutions were plated and plates incubated in 5% CO_2_at 37°C overnight before counting. All assays were performed on three separate occasions in at least four independently inoculated wells.

### Exposure to A549 cells

A549 cells were grown to confluence in half of a 96 well black tissue culture treated plate to confluence in DMEM/F12 with GlutaMAX (Gibco^TM^) plus 10%FCS in 5% CO_2_. No antibiotics were used. The cells remained at confluence for 10 days with q4 day media change. Just before the assay, monolayers were washed twice with PBS. Strain 3735 was grown to an OD_620_ of 0.3, pelleted and the pellet resuspended in RPMI with 1% FCS and 330 μM luciferin. The culture was incubated to allow for uptake of luciferin for 20 minutes at 37°C. The bacterial cells were added to the plate and the plate was spun at 2000 x g for 5 minutes at 4°C. The plate was read and then incubated in 5% CO_2_ at 37°C. The plate was read for luminescence and OD_620_ at 30 and 60 minutes.

### Genome content

Using the PubMLST database, the isolates in the Boston collection were searched via BLAST for full length BlpA, ScbA, ScbB and CSP type [25]. Isolates with either missing BlpA sequence data or typical truncated sizes were marked as BlpA non intact. Any isolate with unique patterns was investigated manually for genome content. The phylogenetic tree was created using the Newick file from Mitchel et al and displayed using iTOL.

## S1. Supplemental methods: Strain construction details

The first reporters made were constructed to have either the *blp* promoter driving a HiBiT tagged version of the bacteriocin BlpI followed by the luciferase gene or the *cibA* promoter region driving a C-terminal HiBiT tagged version of CibA followed by the luciferase gene. These strains were created using a synthetic fragment that created the promoter/ bacteriocin HiBiT fusion and was designed to replace the janus 2 cassette in the transcriptionally silent CEP locus derived from strain 2779 (D39-CEP-Janus2-*luc*). CibAHiBiT was created using a synthetic 992bp fragment created by gBlocks Gene Fragments (IDT) that contains the upstream CEP region, the *cibA* promoter, the CibA ORF with a 10AA linker followed by the HiBiT tag GGDGGGGSGGGGSVSGWRLFKKIS followed by a portion of the luciferase gene in an operon format (sequence listed as primer 5 and 6). The synthetic region was joined to an up and downstream fragment created using primers 1-4. All subsequent fusions were created by amplifying the HiBiT tag/luciferase region of this strain with primers 7 and 8 and fusing via Gibson assembly to a variety of promoters that preceded a peptide of interest. Each peptide carries the same C terminus of a 10AA linker followed by the HiBiT tag. For SPD_145, SPD_0939, SPD_1518, SPD_1745 and SP_1786 region fusions, peptide reporter strains, primers 9-18 were used to place the quorum sensing promoters in front of a luciferase gene in the transcriptionally silent CEP locus by exchanging the constructed fragment into strain 3406 or 3404 which are D39 derivatives with either a disrupted or intact version of the *blpA* gene, respectively and with a janus2 locus in the CEP region. The promoter/peptide regions were all amplified from D39 except ScbA which was cloned from the SP9BS68 genome. Clones were confirmed by PCR and phenotypic responsiveness to respective peptides. Streptococcin B strains were additionally confirmed by sequencing. The P*scbA* reporters were similarly constructed without a HiBiT tagged version of Streptococcin B arranged in an operon format with the luciferase gene from D39 or SP9BS68. The upstream region of the CEP locus was amplified using primers 1 and 2 and joined to the product of 17 and 19 or 17 and 20 to amplify the D39 or SP9BS68 promoters respectively and the downstream product of primers 21 and 4. The HiBiT +/- reporter strains behaved similarly in stimulation assays so the HiBiT tagged version was used to create the ComE binding site deletion. This deletion was created via Gibson assembly using Primers 17 and 22 and 23 and 4 and confirmed by sequencing to carry a deletion of the ComE binding site. The *scbABC* region from SP9BS68 was moved into the D39 strain replacing the native SPD_1786-1784 region as follows: The Janus2 cassette was inserted into the native *scbBC* (SPD_1785-6) region by Gibson assembly using primers 24/26 and 25/27 to create the 5’and 3’ fragment and fusing these to the Janus2 cassette followed by reamplification with primers 28 and 29 transformation into the D39 strains. The cassette was exchanged by transforming with the *scbABC* region of SP9BS68 created with primers 24 and 27. A deletion of the upstream ComR (SPD_1786) and the scbBC region was created by transforming the Janus2 containing strain with a deletion made with primers 30/31 fused to 25/27. A deletion of just the scbBC region was created by transforming the Janus2 containing strain with a deletion made with primers 24/30 joined to 33/25. These strains were confirmed by PCR. The SP18BS74 allelic variant of *comR* was moved into the Janus2 containing strain using genomic DNA from SP18BS74 and selected on sucrose. The construct was back transformed into the Janus2 strain to remove unlinked DNA. The correct insertion was verified using PCR with primer 56 and 19, primer 56 will only anneal to a unique region of the *comR* gene. The *amiCD* deletion was created by moving the *amiCD::janus2* deletion created in Wang et al [3] into the *scbA* reporter strain using primers 34/35. An in-frame, unmarked deletion was created by exchanging this region with the deletion of the *amiCD* region created by Wang et al using the same primers. The *comA*::janus deletion was moved into the Streptococcin B HiBiT expressing stain by transformation with genomic DNA from strain 2898 and selected for on kanamycin and confirmed by PCR.

*cibAB*, *lytA* and *cbpD* deletion strains were created by replacing the entire *cibAB*, *lytA* or *cbpD* ORF with either the spectinomycin (*cibAB*, *lytA*) or kanamycin (*cbpD*) resistance cassette using Gibson assembly. The individual pieces were made with primers 38-55. Fragments were re-amplified and inserted into the indicated strains by transformation. Deletions were confirmed by PCR.

**Figure S1.**
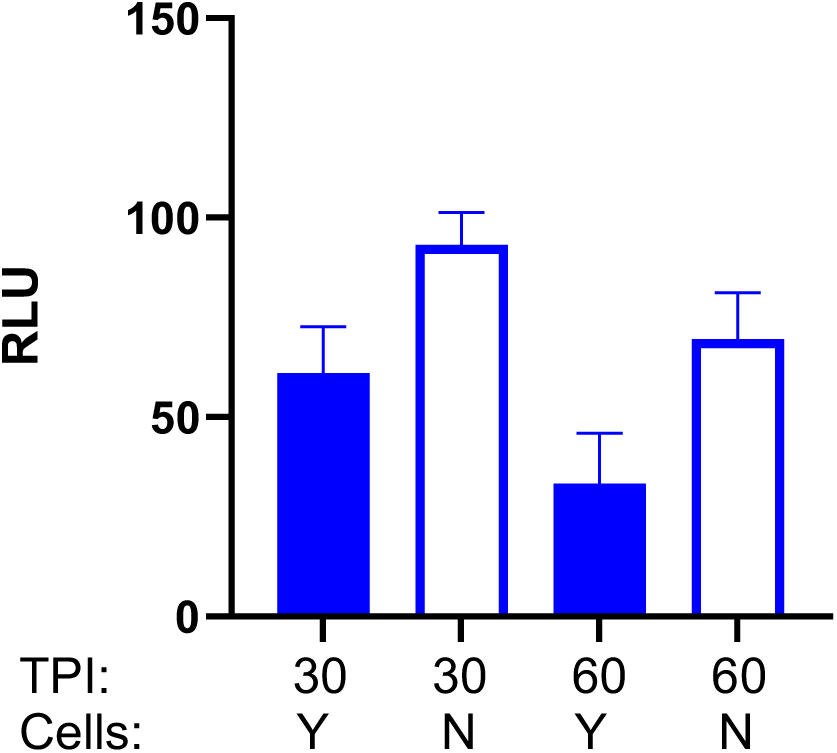
Exposure to live A549 cells does not induce the D39 scbBC promoter. Reporter strain 3735 with the D39 *scbBC* promoter driving luciferase production was incubated with live A549 cells at 5% CO2 in RPMI + 1%FCS for the indicated times in the presence of luciferin. RLU was determined in wells with and without cells. TPI= Time post infection. Assay was done with 6 replicates for each condition and was repeated once for verification. A representative assay is shown.

**Table S1.**
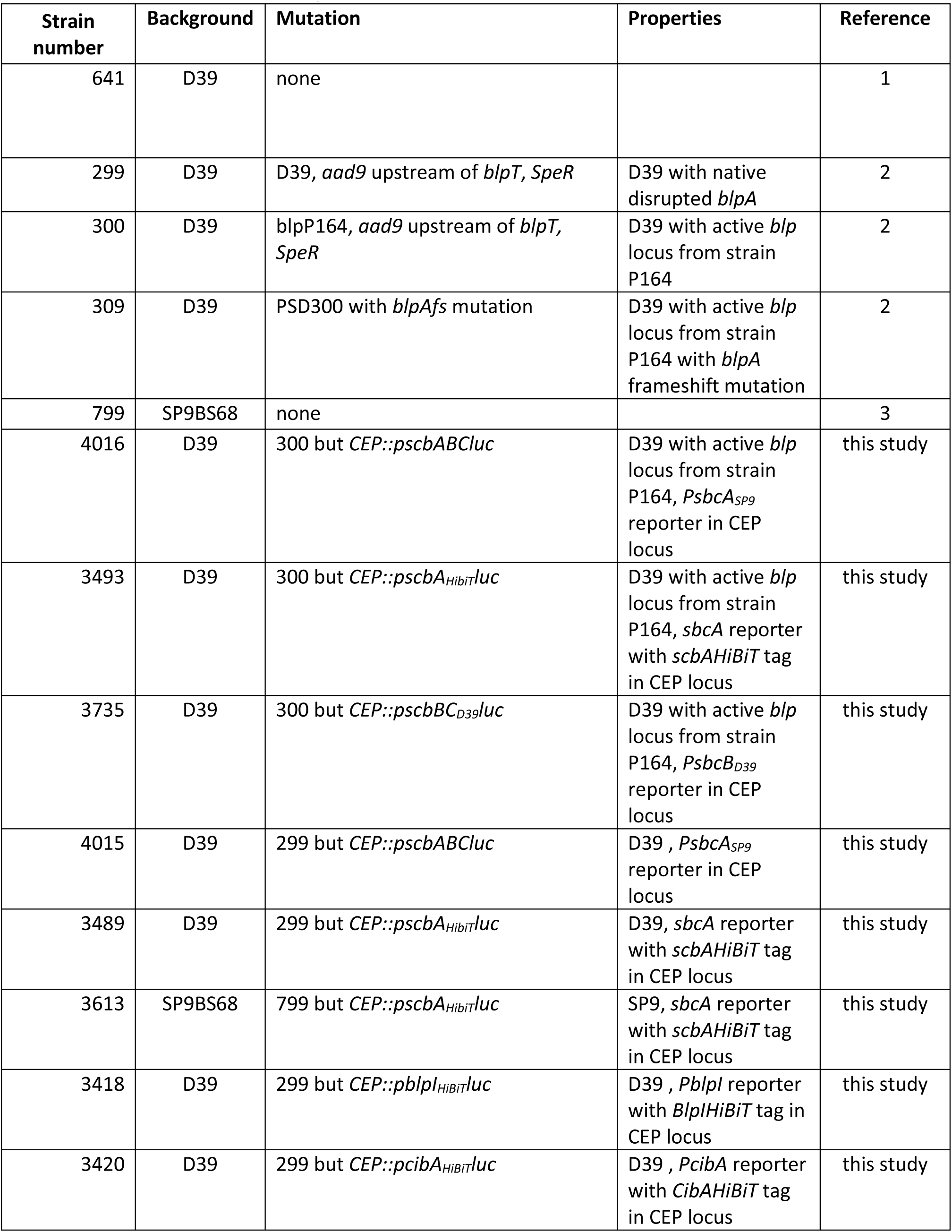
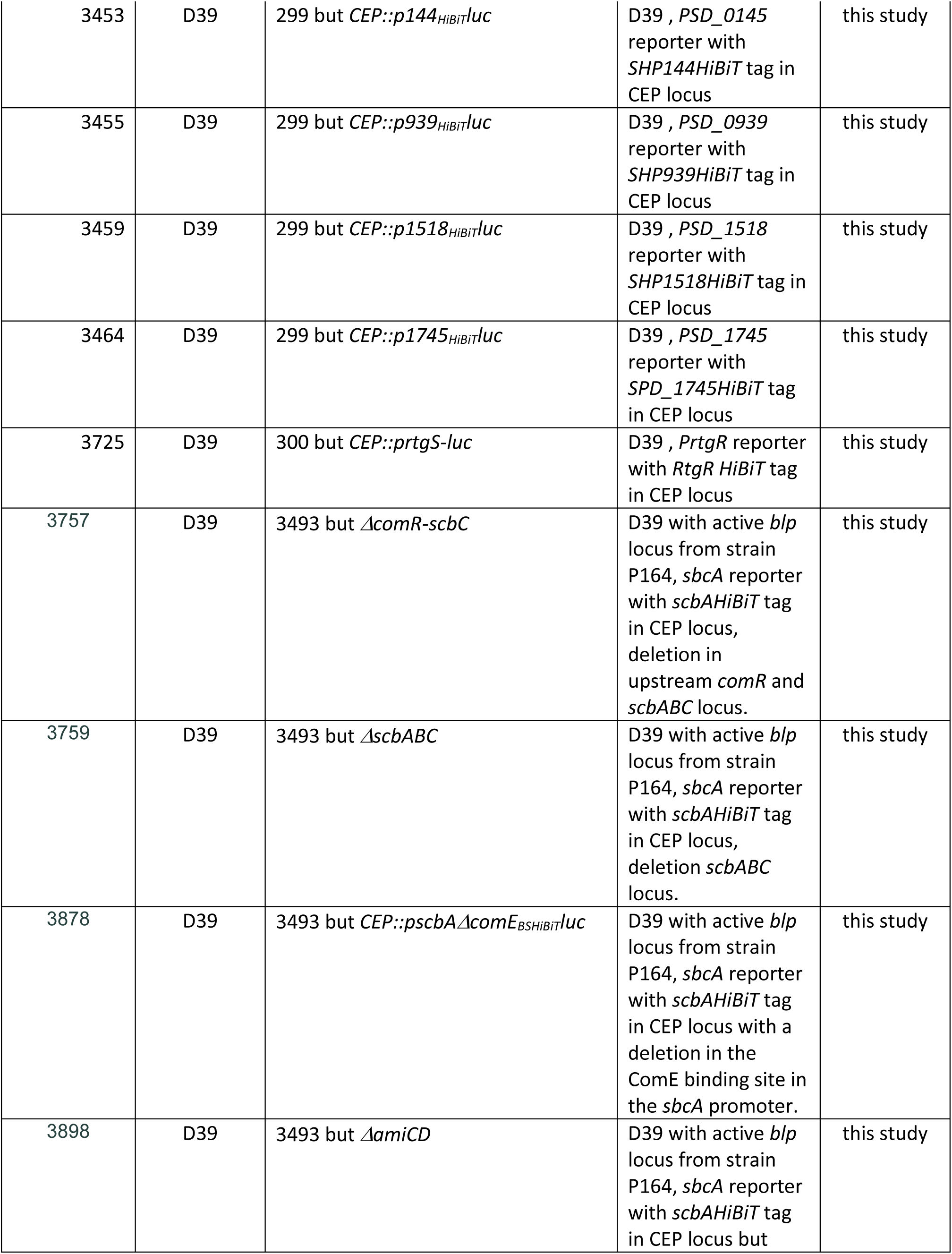
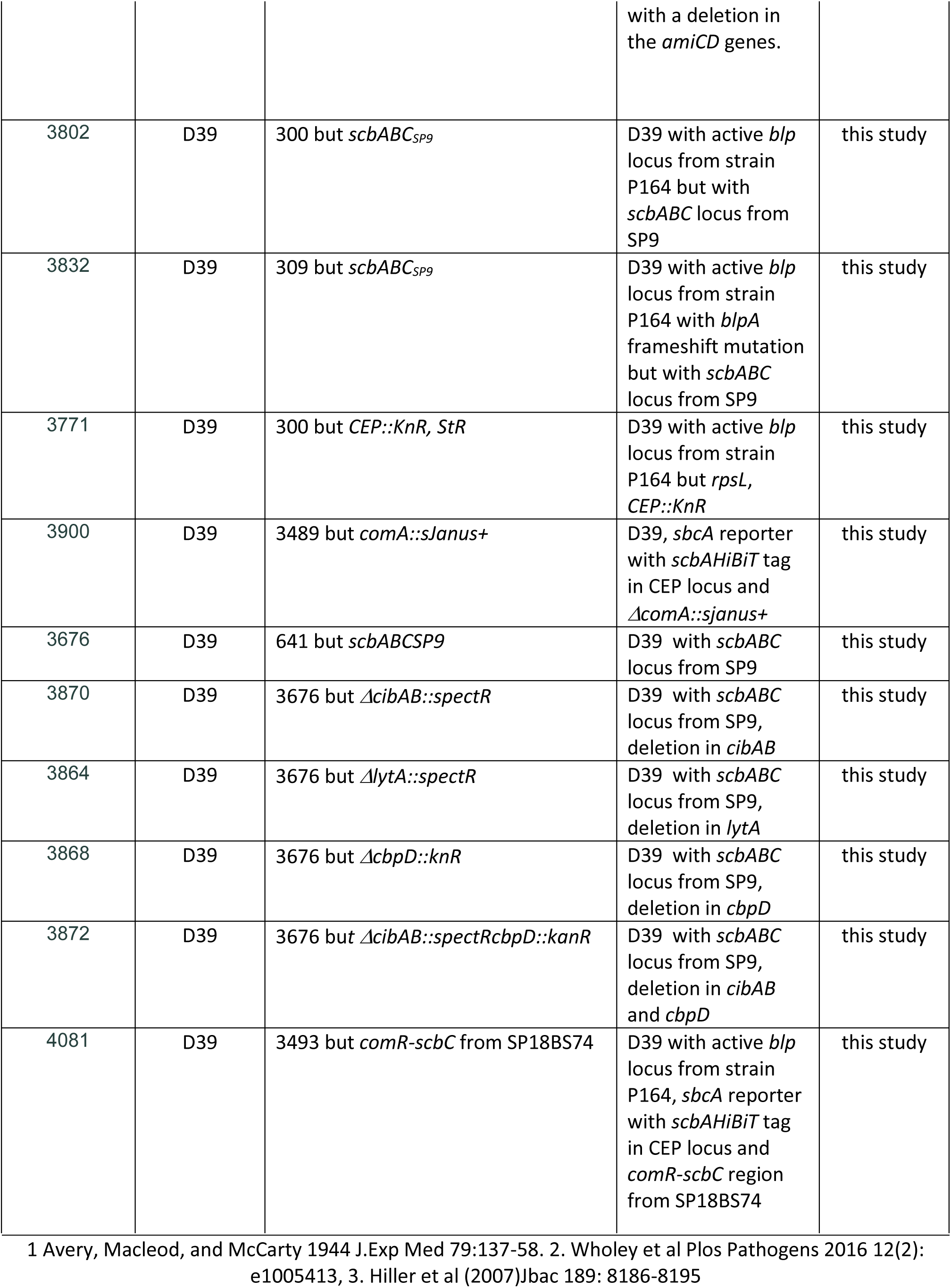
Strains used in this study.

**Table S2.**
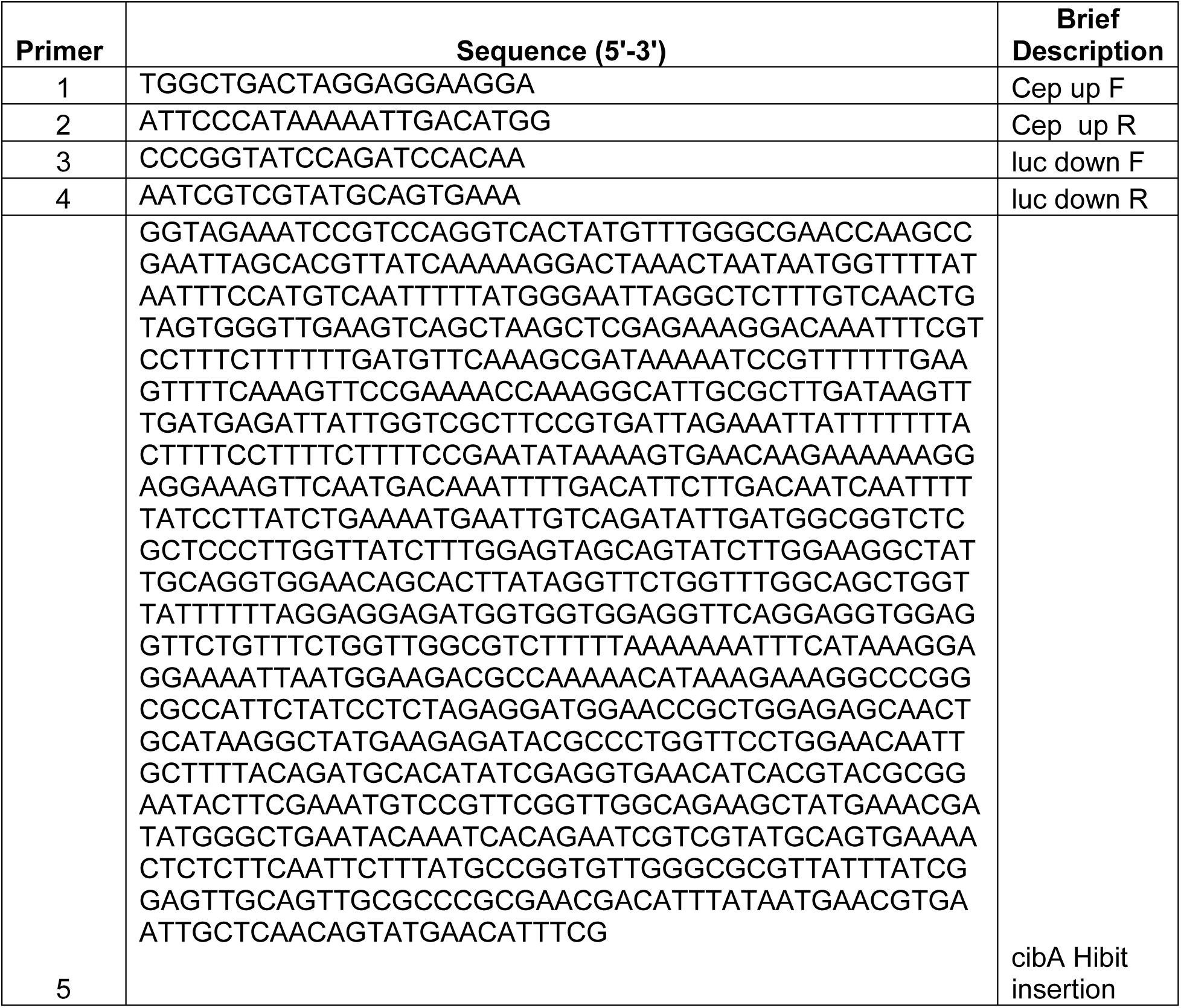
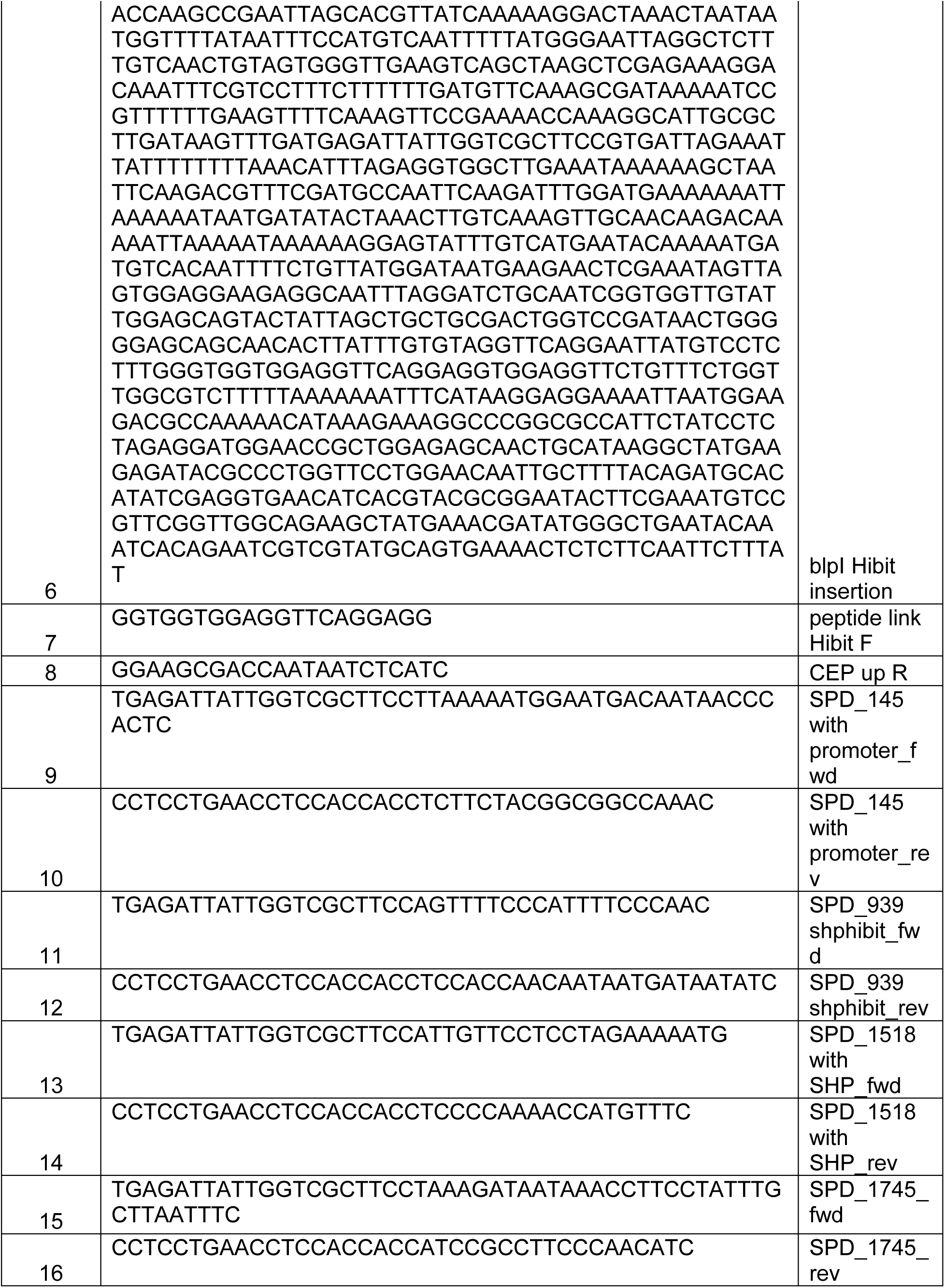
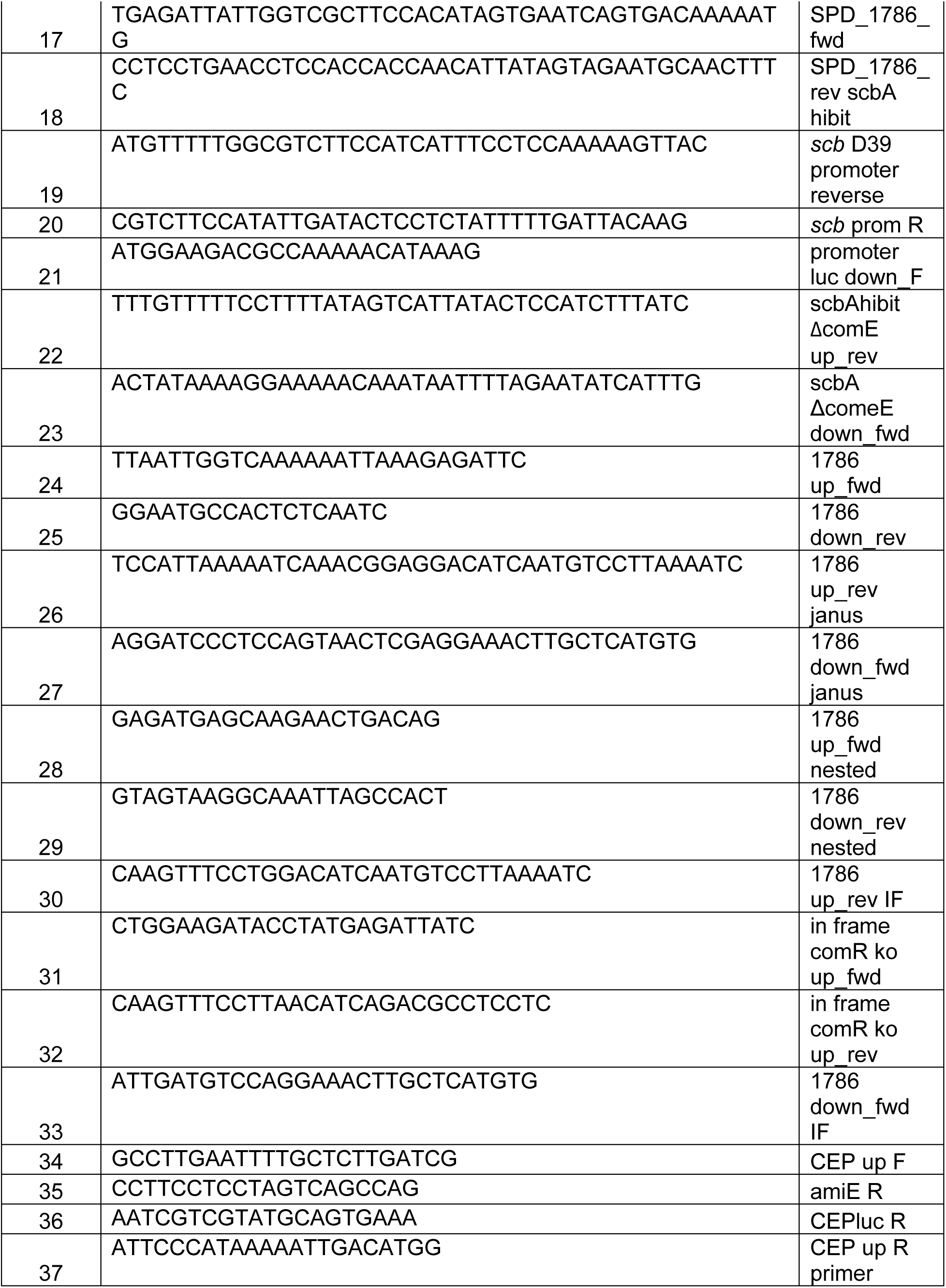
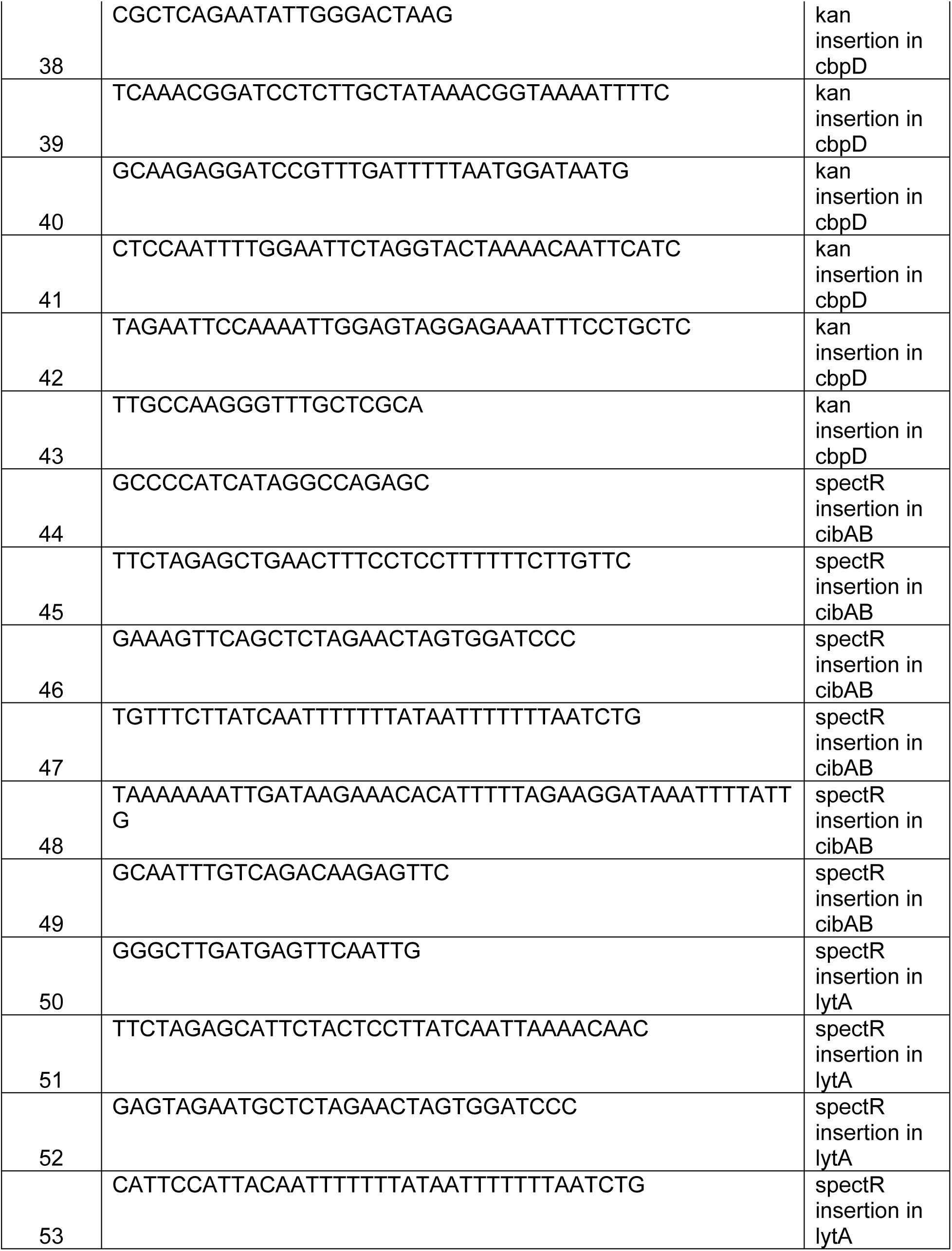
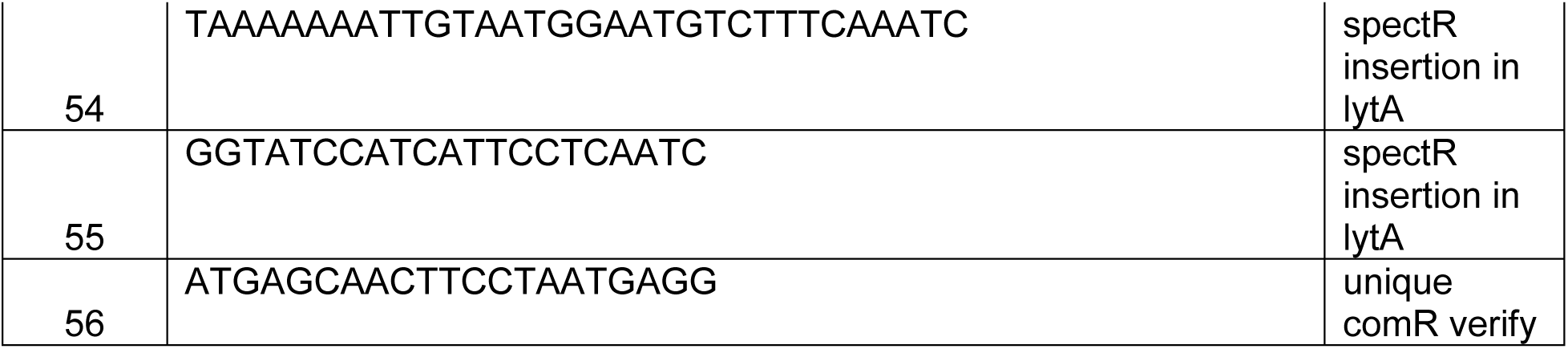
Primers/ synthetic fragments used in this study.

**Table S3.** Peptides used in this study.

| Peptides | Sequence |
| --- | --- |
| CSP1 | ENRKSJFFRDFUKQRKK |
| BlpC2 | GWWEELLHETILSKFKITKALELPIQL |
| SHP144 | EWVIVIPFLTNL |
| SHP939 | DIIIVGG |
| SHP1518 | LIWFETWFWG |
| PhrA | SNGLDVGKAD |
| *S.pseudopneumoniae* XIP | GDWWHG |
| RtgS | AIIFPWGWP |

